# Insilico analysis of structural and functional impact of SNPs in Pleckstrin

**DOI:** 10.1101/782623

**Authors:** Arpit Kumar Pradhan, Ashwin Jainarayanan, M. A. Nithishwer, Shyamasree Ghosh

## Abstract

Pleckstrin (PLEK) gene has been associated with a variety of disorders including autoimmune, inflammatory diseases and cancer. Mutation in this gene has been reported to be associated with autoimmune celiac disease (CD), increased atrophy in multiple sclerosis (MS), obstructive sleep apnea (OSA), abdominal aortic aneurysms, over expression in inflammatory disorders including periodontis, risk for ependymoma relapse, bladder cancer, melanoma, lung, and colorectal cancer revealing the importance of study of the PLEK. PLEK gene has been reported from other animals and therefore we have studied the molecular evolution of the PLEK gene by *insilico* approaches. Single nucleotide polymorphisms (SNPs), in humans have been reported to cause potential structure-function alteration in proteins. In this study we have tried to understand by *insilico* approaches the (i) molecular evolution of PLEK and (ii) the impact of potentially deleterious single non-synonymous SNPs (nsSNPs) on the structure and function of Pleckstrin protein. We report for the first time using molecular dynamic simulation (MDS), the impact of SNPrs17035364 and rs3816281 on the structural alterations of Pleckstrin with implications in altering its biological function which may find importance as diagnostic markers.

## 1. Introduction

Pleckstrin (PLEK) gene has been reported to be associated in a number of diseases including autoimmune multifactorial celiac disease (CD), triggered by gluten ingestion^1^, rheumatoid arthritis (RA), multiple sclerosis (MS), cardiovascular disease (CVD), inflammatory disorders, ulcerative colitis (UC), periodontitits^2^, obstructive sleep apnea (OSA), abdominal aortic aneurysms, increased risk for ependymoma relapse and cancer including bladder, melanoma, lung and colorectal cancer. For an in depth review we refer the readers to (Rebecchi and Scarlata 1998)^3^ In mice, the PLEK gene is reported to be upregulated in transformation-resistant cells^4^. PLEK gene has been reported from different animals and considering the importance of the PLEK gene and its role in humans and mice as animals models we have tried to understand the molecular evolution of the PLEK gene amongst different animals by *insilico* approaches.

Understanding the structure and function of the PLEK protein finds importance in understanding the biology of diseases in which they play major role PLEK protein. Human, PLEK protein has two pleckstrin homology(PH) domains that plays a major biological role as a substrate of protein kinase C in platelets^5^. This domain can bind phosphatidylinositol lipids within biological membranes including phosphatidylinositol (3,4,5)-trisphosphate and phosphatidylinositol (4,5)-bisphosphate), and the βγ-subunits of heterotrimeric G proteins and protein kinase C^5^.

Non-synonymous SNPs (nsSNPs) results in amino acid sequence alterations in proteins and can lead to structural and functional alteration in the protein structure^(6–7)^. Advancements in high throughput sequencing have increased the number of DNA sequence variants being submitted but the real challenge remains in understanding their role in diseases. Therefore, an efficient as well as effective method is necessary to screen the most deleterious variant that leads to an alteration in structure-function activity of the protein.

Computational models and tools have helped screening of SNPs that affect the function of proteins and provides an insight to the structural basis of disease-causing mutation. *Insilico* approaches to study SNPs have enabled us to understand the structural modifications in certain protein and their direct impact in gall bladder cancer^6, 7^.

Since mutations in the PLEK gene has been known to be associated with a variety of disorders, and reported from different animal systems, in this study we have performed by *insilico* approaches to understand the (i) molecular evolution of PLEK and (ii) the impact of single nucleotide polymorphisms (SNPs) on the structure and function of Pleckstrin gene.

## 2. Materials and methods

### 2.1. PLEK Sequence retrieval and alignment

The PLEK sequences were retrieved from the NCBI database. The PLEK sequence was submitted to the blast server and was searched for different taxa. Alignment of sequences obtained from different organism was done using the ClustalW^8^ and MUSCLE^9^ alignment.

### 2.2. Calculation of evolutionary time

In order to explore the molecular evolution of, amino acid sequences for different classes of animals were compared. The number of changes of amino acids per 100 amino acids was calculated by comparing mammals with birds, birds with amphibians and mammals with amphibians. Human PLEK sequence is considered the most recent and hence the evolutionary time of human sequence is considered as zero million years. The average changes were calculated and the radiation of mammalian PLEK was plotted against million years. For calculating the radiation of non-human primates, human sequence was compared with those of Gorilla, Common Chimpanzee, Sumatran orang-utan, Bonobo, Rhesus macaque, Olive baboon, Crab-eating macaque, Green monkey, Drill, Sooty mangabey, Southern pig-tailed macaque, Northern white-cheeked gibbon, Nancy Ma’s night monkey, Angola colobus and white-headed capuchin. For calculating total mammal radiation, Leopard, Cheetah, Siberian Tiger, vesper bat, Chinese rufous horseshoe bat and Tupaia sequences were compared with human PLEK sequences. In these cases, the average value representing the amino acid change/100 amino acids were considered. For calculation of birds with amphibians, African clawed frog sequence was compared with Grey crowned crane, Peregrine falcon, Rock dove, Chicken and Mallard sequences. Amphibians and mammals were analyzed by using PLEK from Wild Bactrian camel, Goat and Giant Panda with that of the African clawed frog. Similarly, amphibians and fishes were compared by comparing the sequence of the African clawed frog with Zebrafish, Clownfish, and Black striped Livebearer. For a similar analysis, histone-4 was used as a highly conserved protein and Cytochrome-C as a semi-conserved protein^10^.

### 2.3. Selection of SNPs for insilico analysis

The dbSNP database, being the most extensive database, was availed to procure the SNPs in the PLEK sequence in spite of its limitations of containing both validated and nonvalidatedpolymorphisms^11^. The Global Minor Allele Frequency (MAF) was taken into consideration while choosing the SNPs for our analysis. All the missense SNPs were listed and were selected for the further analysis.

### 2.4. Prediction of deleterious nsSNPs by insilico tools

Missense SNPs leads to amino acid variations. PolyPhen-2 and SIFT were used to predict the damaging effect of missense SNPs on structure and function of the protein by analysis of multiple sequence alignment and protein three-dimensional (3D) structure, the protein sequence, the position at which substitution takes place, the amino acid being substituted, and the amino acid present in the variant type^12,13^. In order to study the functional implications of the SNPs, each SNP with their respective rsIDs was uploaded, and the study was done for every nsSNPs. The outcome of the SNP prediction could be categorized as probably damaging, possibly damaging, or benign according to the PolyPhen-2 score ranging from 0 to 1. The score refers to the amino acid substitution in the variant type being damaging. These scores are HumDiv scores, compiled from all damaging alleles with known effects on molecular function, and HumVar scores which consist of human disease-causing mutations. The closer the HumDiv score is to 1, it is indicative of greater damaging nature of the SNP.

SIFT takes in a query sequence (in our case the rs ID was the input) and predicts the tolerated or deleterious substitution based on the information of multiple alignments for every position of the query sequence^13^. SIFT then obtains the multiple alignment of sequences and calculates the normalized probabilities for each substitution. Substitutions at each position could fall into two categories, either damaging or tolerated based on whether the normalized probabilities are less than or greater than a tolerant index of 0.05. The missense SNPs were analyzed to obtain the damaging SNPs and these were analyzed further for their structural and functional implication in PLEK.

### 2.5. Protein structure prediction

I-TASSER webserver^14^ was used for the generation of the 3D structure of the protein and prediction of the biological functions of protein molecules from amino acid sequence. It provides five models based on the amino acid sequence and each model is assigned an individual C-score (confidence scores) which is calculated on the basis of significance of threading template alignments and the convergence parameters of the structure assembly simulations. A higher C-score indicates greater confidence and vice-versa. It also provides a TM (template modeling) Score which is a measure of the structural similarity between two structures. A TM-score >0.5 indicates a model of correct topology whereas a TM-score <0.17 indicates a random similarity. It also predicts solvent accessibility and ligand binding sites. Protein sequence was submitted to the webserver and the protein structure was obtained.

### 2.6. Modeling of the mutant protein structure

The mutated model for the PLEK protein corresponding to the respective amino acid substitution was generated by Swiss-Pdb Viewer (v4.10)^15^. The amino acid in the native sequence is replaced with the variant and the .pdb files were saved for further analysis.

### 2.7. MD simulations in water

The Desmond package^16^ from Schrodinger 2017-2 was used to run the molecular dynamic simulation^17^. Predefined TIP3P water model was used for the simulation of water molecules. Orthorhombic periodic boundary conditions buffered at 10 Å distances were set up to specify the shape and size of the repeating unit. To neutralize the system, appropriate counter ions (Na^+^/Cl^-^) were added and were placed randomly in the solvated system. The minimization and relaxation of the protein/protein-ligand complex was performed after setting up the solvated system by NPT ensemble using default protocol of Desmond as followed elsewhere; which includes a total of 8 stages which includes series of minimization and short simulation steps^18–19^.

*Summary of Desmond’s MD simulation stages*:

Stage 1—task (recognizing the simulation setup parameters)
Stage 2—simulate, Brownian Dynamics NVT, T = 10 K, small timesteps, and restraints on solute heavy atoms, 100ps
Stage 3—simulate, NVT, T = 10 K, small timesteps, and restraints on solute heavy atoms, 12ps
Stage 4—simulate, NPT, T = 10 K, and restraints on solute heavy atoms, 12ps Stage 5—solvate_pocket
Stage 6—simulate, NPT and restraints on solute heavy atoms, 12ps Stage 7—simulate, NPT and no restraints, 24ps Stage 8—simulate

All the molecular dynamic simulation was performed with the periodic boundary condition in the NPT ensemble using OPLS force field parameter. The temperature and pressure were set at 300K and 1 atmospheric pressure respectively using noosehover temperature coupling and isotropic scaling. The recording interval was set at 48ps and approximate number of frames at 250.

### 2.8. Analysis of molecular dynamics trajectory

The Molecular Dynamics Simulation trajectories were analyzed using simulation event analysis and simulation interaction diagram programs available with the Desmond Module. The Root mean squared deviation (RMSD) and Root mean squared Fluctuation (RMSF) data for Wild-type and mutated proteins were calculated from the simulation interaction diagram. The total Energy and Coulombs Energy was obtained from the simulation files whereas the number of intra-molecular Hydrogen bonds and Radius of gyration was obtained from the Simulation event analysis. The data obtained was then graphed using R project for Statistical Computing. SEA is used towards analyzing each frame of simulated trajectory output while SID is utilized to analyze parameters during the simulation time.

## 3. Results

### 3.1. Evolution of PLEK gene

Molecular evolution of PLEK was analyzed in detail, thereby extracting partial or full length PLEK sequences from various databases.

Further, evolutionary time of PLEK was computed by calculating the number of amino acid changes per 100 amino acids as mentioned previously. The analysis performed indicates that the PLEK gene has evolved during the transition from Silurian to Devonian era (approximately 400 mya). The evolutionary slope of PLEK suggests that it has undergone different magnitudes of selection pressure during different periods of time (Fig 1). The variation was low during the radiation of mammals and lower during the radiation of non-human primates. In the Jurassic and Cretaceous era, PLEK has undergone a few changes which indicate a process of molecular stabilization coinciding with the radiation of mammals.

**Fig 1:**
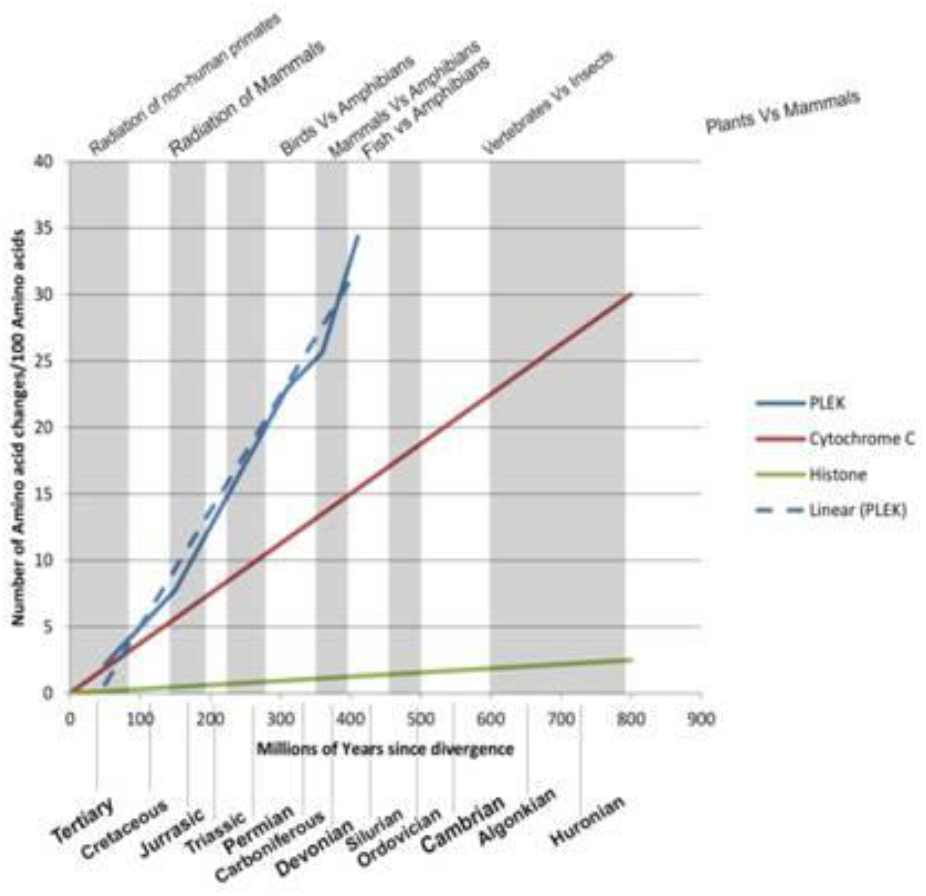
Conservation analysis performed on PLEK in comparison to Cytochrome C (a semi-conservative protein) and Histone H4 (a conserved protein). The values for histone-H4 and Cytochrome-C are plotted as best-fit while values for PLEK are plotted according to the original values.

### 3.2. Selection of SNPs for PLEK gene

The protein sequence for the PLEK gene NCBI Reference Sequence: NP_002655.2 and related SNPs were obtained from dbSNP(http://www.ncbi.nlm.nih.gov/SNP/) and were subjected to various computational analyses. Out of several missense SNPs, the SNP having a higher minor allele frequency (MAF (>0. 05)) were considered for further analysis. MAF is linked to statistical power of the study i.e. there exists an inverse relationship between MAF and sample size to detect variant allele of a SNP in a given sample population. Therefore, SNPs with a MAF of 0.05 or more are generally targeted in the majority of large-scale genome studies.

### 3.3. Analysis of mutations in nsSNPs

The missense SNPs were submitted to the polyphen server. Two SNPs (rs3816281 and rs17035364) were predicted to be damaging by Polyphen. rs3816281 was predicted to be damaging with a humdiv score of 0.997 and humvar score of 0.935. rs17035364 had a humdiv score of 0.990 and humvar score of 0.674. rs17035364was also predicted to be damaging by SIFT.

Native sequence was submitted to I-Tasser in their respective FASTA format. Five models were generated, out of which the first model which had a higher C–Score of – 1.4 and a TM score of 0.54±0.15 was considered for the analysis.

### 3.4. Molecular Dynamics (MD Simulations)

To make a comparative study the conformational changes in the proteins due to the mutations, molecular dynamics (MD) simulation was carried out for each protein^17^. Parameters that were analyzed include radius of gyration, coulomb’s energy, total energy, total number of intra-molecular hydrogen bonds, RMSD (Root mean squared deviation), RMSF (Root mean squared fluctuations) as a time dependent function of MDS. The data obtained was plotted using R statistical tool and a comparative study was made. The time duration of the simulation used is optimal for the protein’s native state and to facilitate various conformations. Further, recent studies have shown that the dynamics of a single protein molecule are self similar and resemble the same, irrespective of duration.

In order to understand the effects of mutations on the structure of the protein, the RMSD values of the wild type/native and other mutant proteins were analyzed. The RMSD of the protein backbone were calculated during the simulation with reference to its initial structure. RMSD of Wild type/native was found to be relatively more stable than the other two mutated proteins carrying rs3816281 and rs17035364 (Fig 2) respectively. The RMSD of the wild type had a mean value of 5.4 Å, whereas the mutant proteins had a higher mean (rs3816281 had a mean value of 6.51 and rs17035364 had a mean value of 5.79). Among all residues, the highest fluctuation was found approximately at residue number 300. Fig. 3 clearly demonstrates the destabilizing effects of the SNPs on proteins.

**Fig 2:**
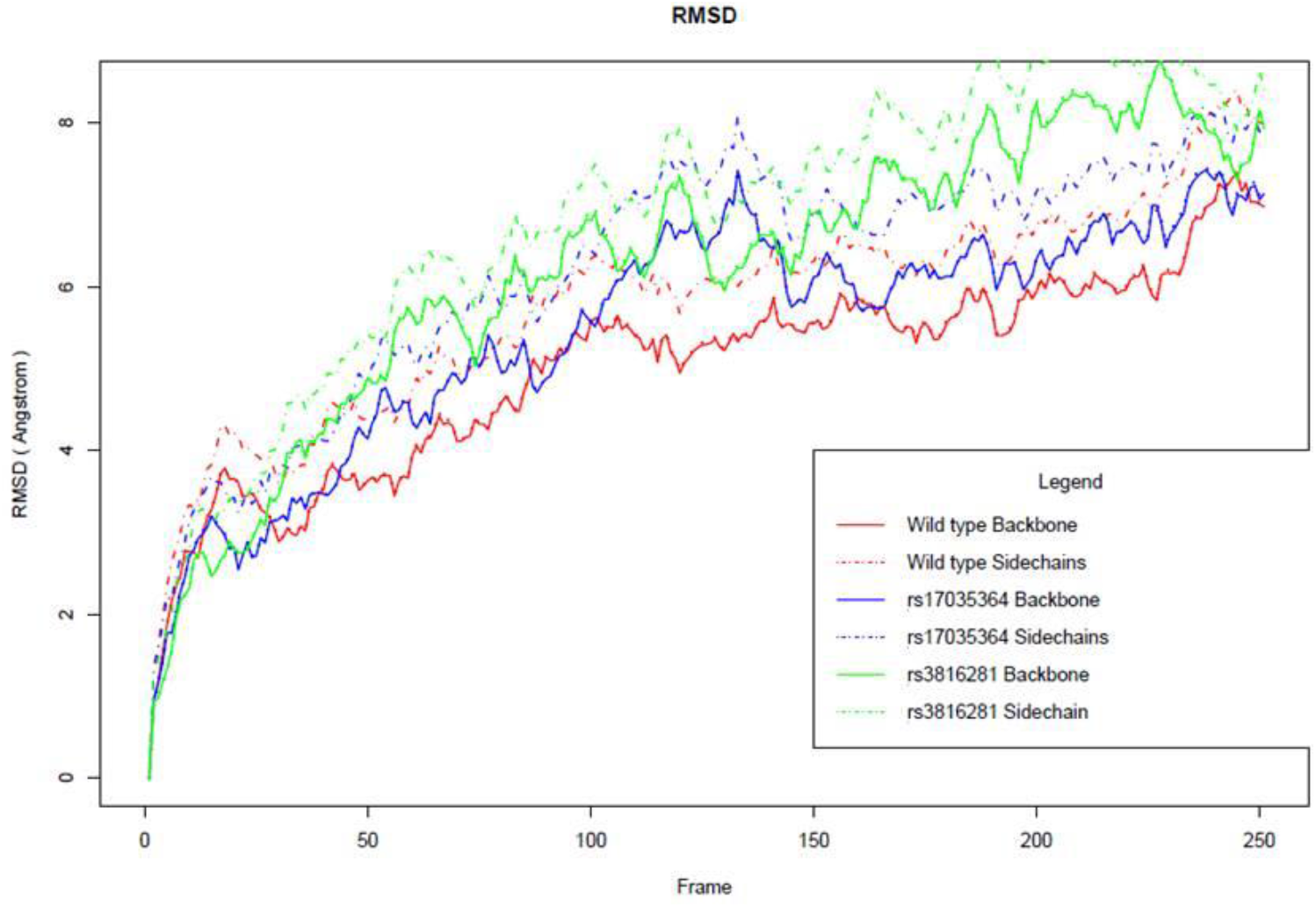
RMSD values of the wild type protein along with those of the associated mutant proteins.

**Fig 3:**
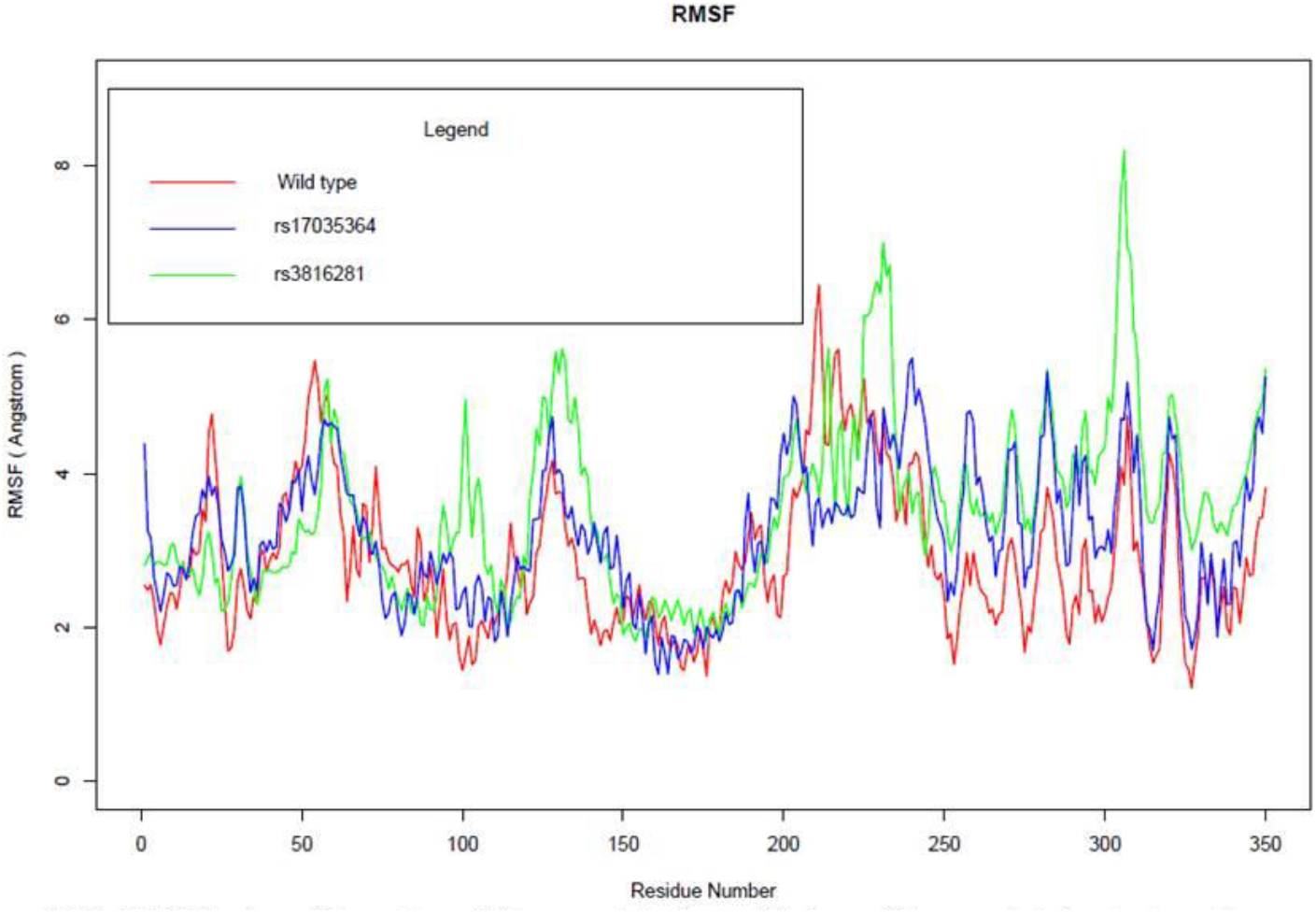
RMSF values of the native wild type protein along with those of the associated mutant protein.

The RMSF of each residue was also monitored to determine the effects of mutation on the protein residues’ dynamic behavior. The results strongly suggeststhat residue level fluctuations for rs3816281 mutation is high with respect to both the native and the rs17035364 (Fig 3). From the analysis it is evident that rs3816281 mutated protein has the highest degree of flexibility.

Analysing the energy parameters for the MDS trajectories revealed that the total energy of wild type/native ranged between −289040.6 Kcal/mol and −288823.6 Kcal/mol. Whereas, the range was found to be significantly higher (−288743.3 Kcal/mol to −288506.8 Kcal/mol) for rs17035364 and significantly lower (−289315.9 Kcal/mol to −289056.4 Kcal/mol) for rs3816281. As compared to the wild type, SNPs rs3816281 has the lower energy whereas mutation rs17035364 has a higher energy (Fig 4).

**Fig 4:**
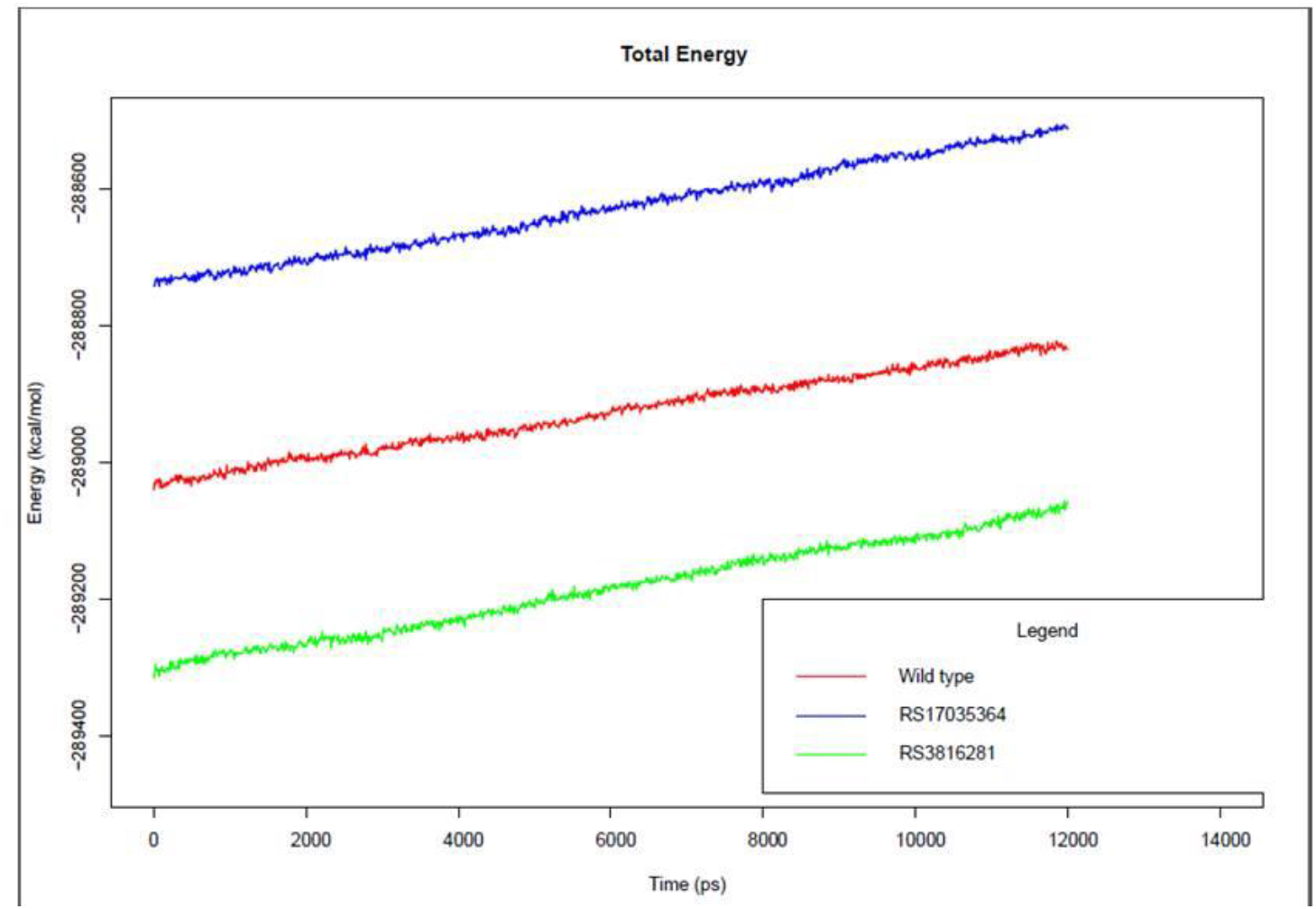
Total energy of the PLEK protein compared to that of each mutant protein.

Study of the total numbers of intra-molecular Hydrogen bonds of the wild type and mutant proteins (Table 1) revealed that the highest number of number of Hydrogen bonds occurred in the wild type followed by rs17035364 and the lowest being rs3816281. On the other hand, rs3816281 had the largestrange. This indicates that rs3816281 has greater flexibility compared to either the wild type/native or rs17035364 forms (Fig 5) confirmed from the RMSF data.

**Fig 5:**
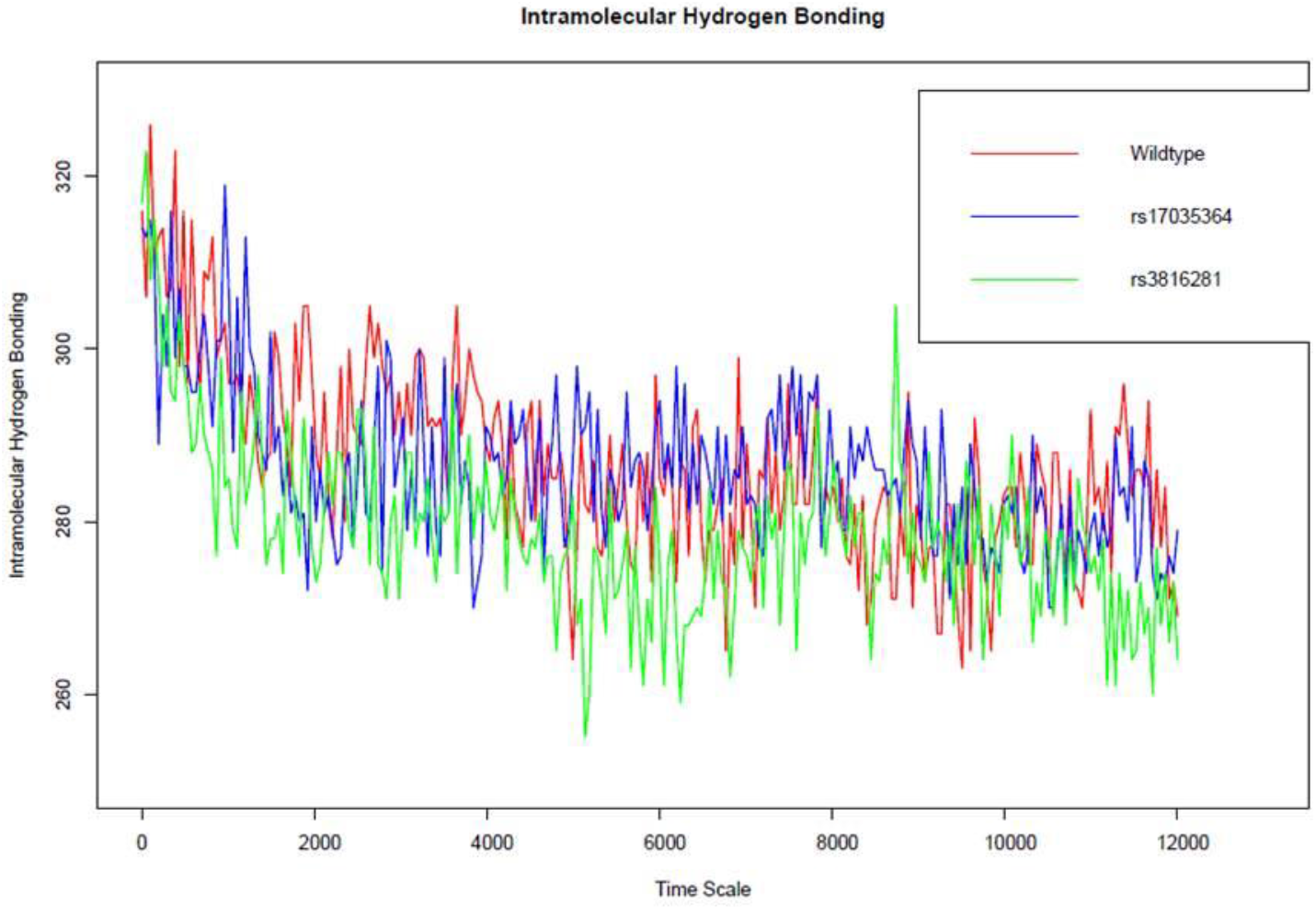
Number of intramolecular Hydrogen bonds of the PLEK protein compared to that of each mutant protein.

**Table 1.**
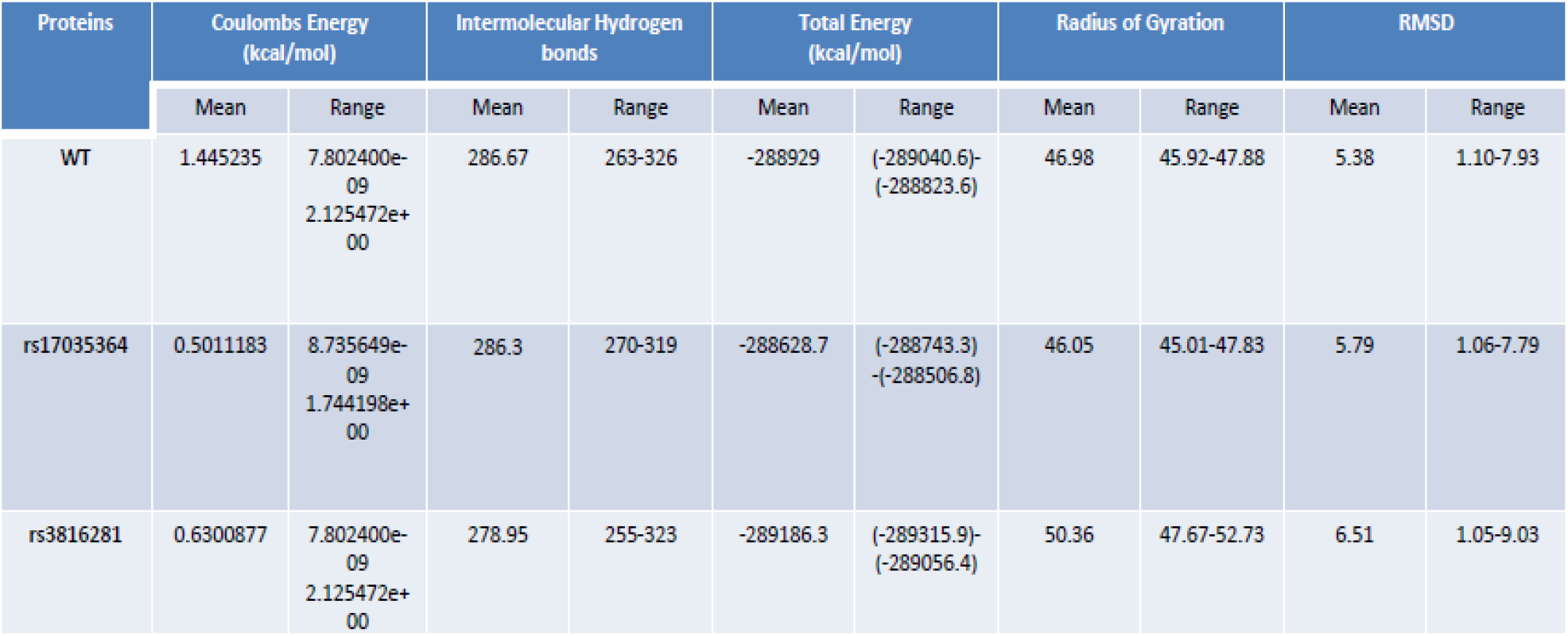
Statistical data from the MD simulation trajectories of wild type/native and mutated PLEK proteins.

| Proteins | Coulombs Energy (kcal/mol) |  | Intermolecular Hydrogen bonds |  | Total Energy (kcal/mol) |  | Radius of Gyration |  | RMSD |  |
| --- | --- | --- | --- | --- | --- | --- | --- | --- | --- | --- |
|  | Mean | Range | Mean | Range | Mean | Range | Mean | Range | Mean | Range |
| WT | 1.445235 | 7.802400e-09<br>2.125472e+00 | 286.67 | 263-326 | -288929 | (-289040.6)-<br>(-288823.6) | 46.98 | 45.92-47.88 | 5.38 | 1.10-7.93 |
| rs17035364 | 0.5011183 | 8.735649e-09<br>1.744198e+00 | 286.3 | 270-319 | -288628.7 | (-288743.3)-<br>(-288506.8) | 46.05 | 45.01-47.83 | 5.79 | 1.06-7.79 |
| rs3816281 | 0.6300877 | 7.802400e-09<br>2.125472e+00 | 278.95 | 255-323 | -289186.3 | (-289315.9)-<br>(-289056.4) | 50.36 | 47.67-52.73 | 6.51 | 1.05-9.03 |

The radius of gyration of the wildtype/native and mutant proteins were analyzed to measure their compactness (Fig 6). From Table 1, it is revealed that rs17035364 mutation possessing protein has the highest compactness with a mean radius of gyration of 46.05 A, whereas the rs3816281 mutation containing protein has the least compactness.

**Fig 6:**
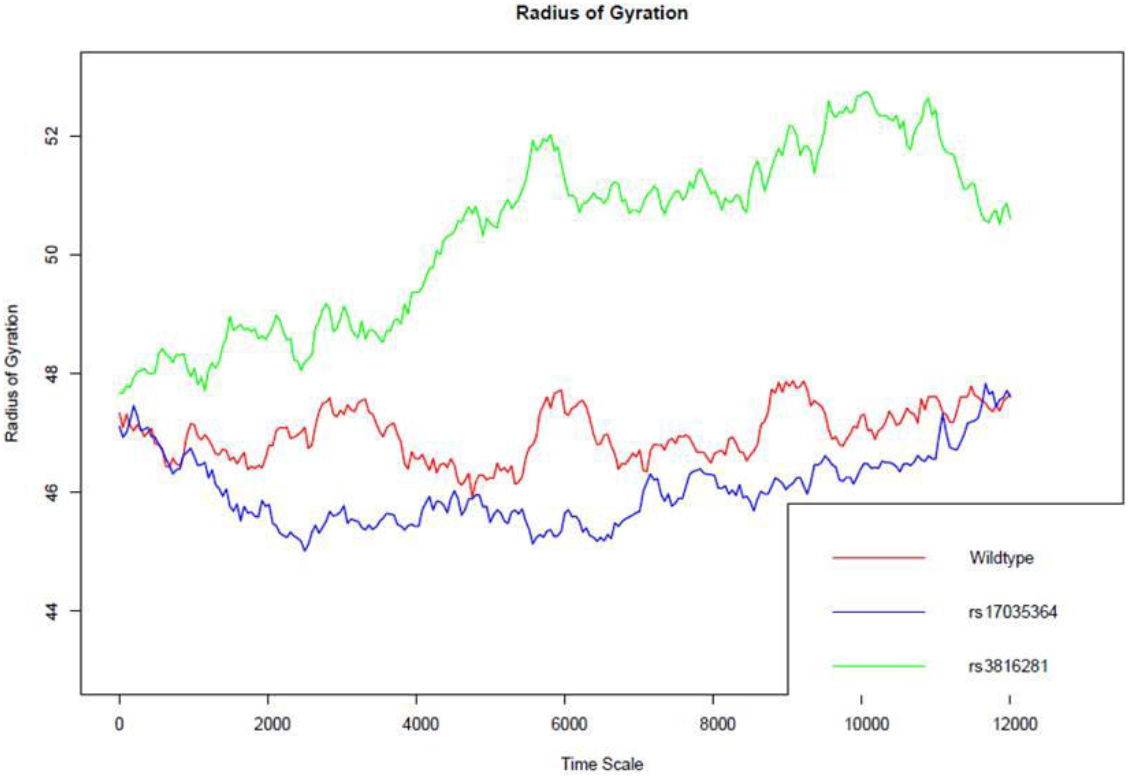
Radius of Gyrations of the PLEK protein compared to that of each mutant protein.

The structural changes in both the SNPs were captured at different time points. Fig 7–10). We recorded snapshot of the PLEK mutant trajectory for every nanosecond of the MD simulation forrs17035364 and rs3816281 (Fig. 7, 9) and superimposed images of pre (Red) and post (Green) MD structures for visualization of the rs17035364 andrs3816281 (Fig. 8, 10) revealing distinct differences.

**Fig 7:**
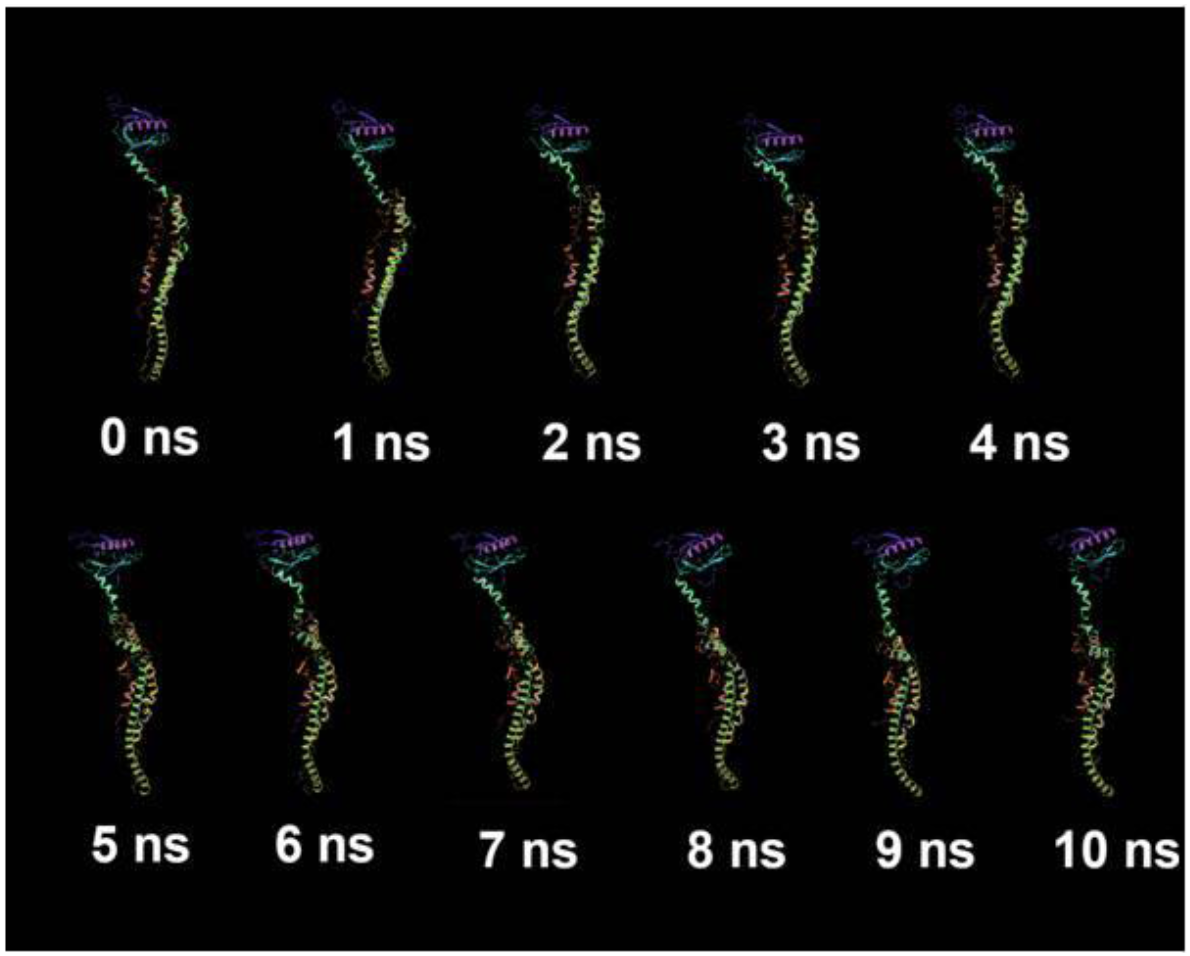
Snapshot of the rs17035364 PLEK mutant trajectory for every ns of the MD simulation.

**Fig 8:**
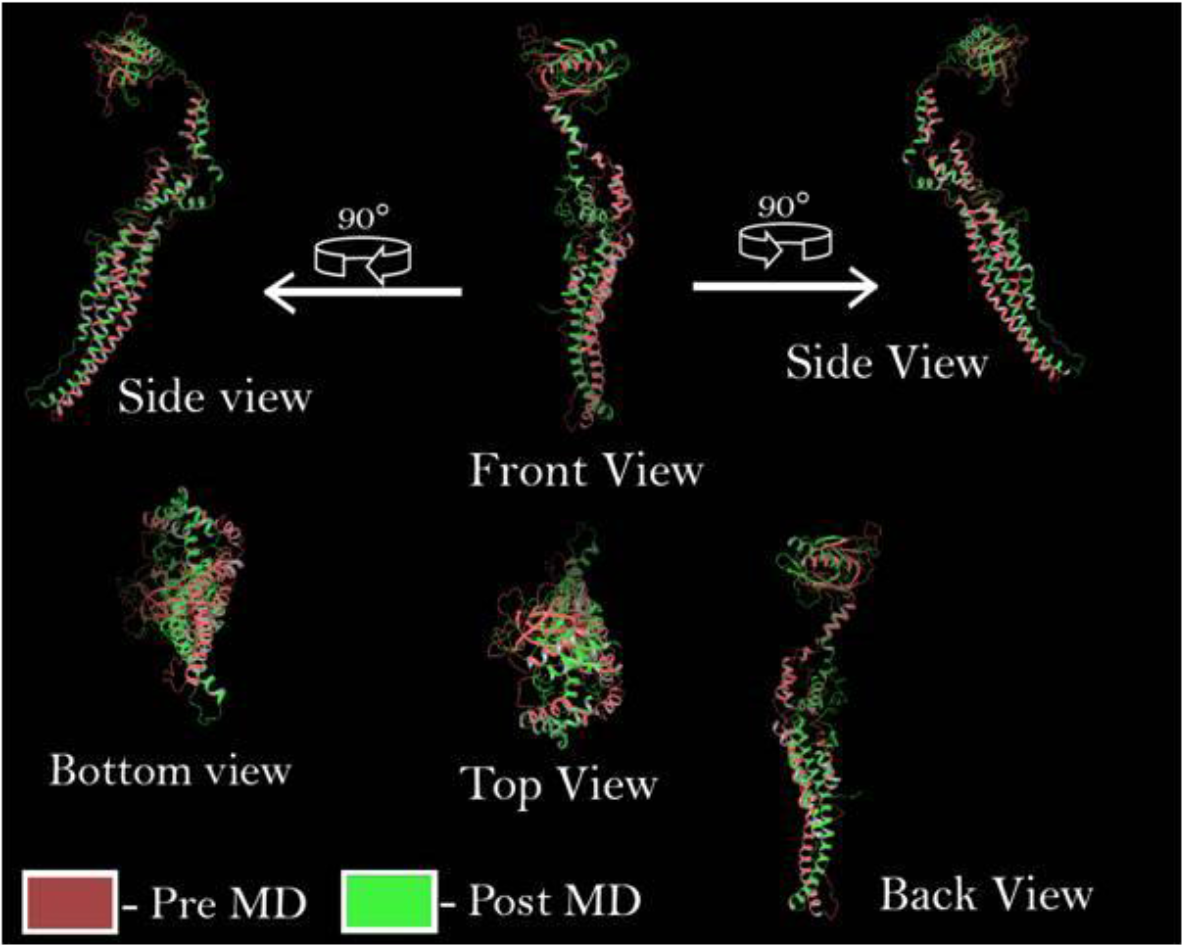
Visualization of the rs17035364 mutant PLEK protein superimposed of pre (Red) and post (Green) MD structures

**Fig 9:**
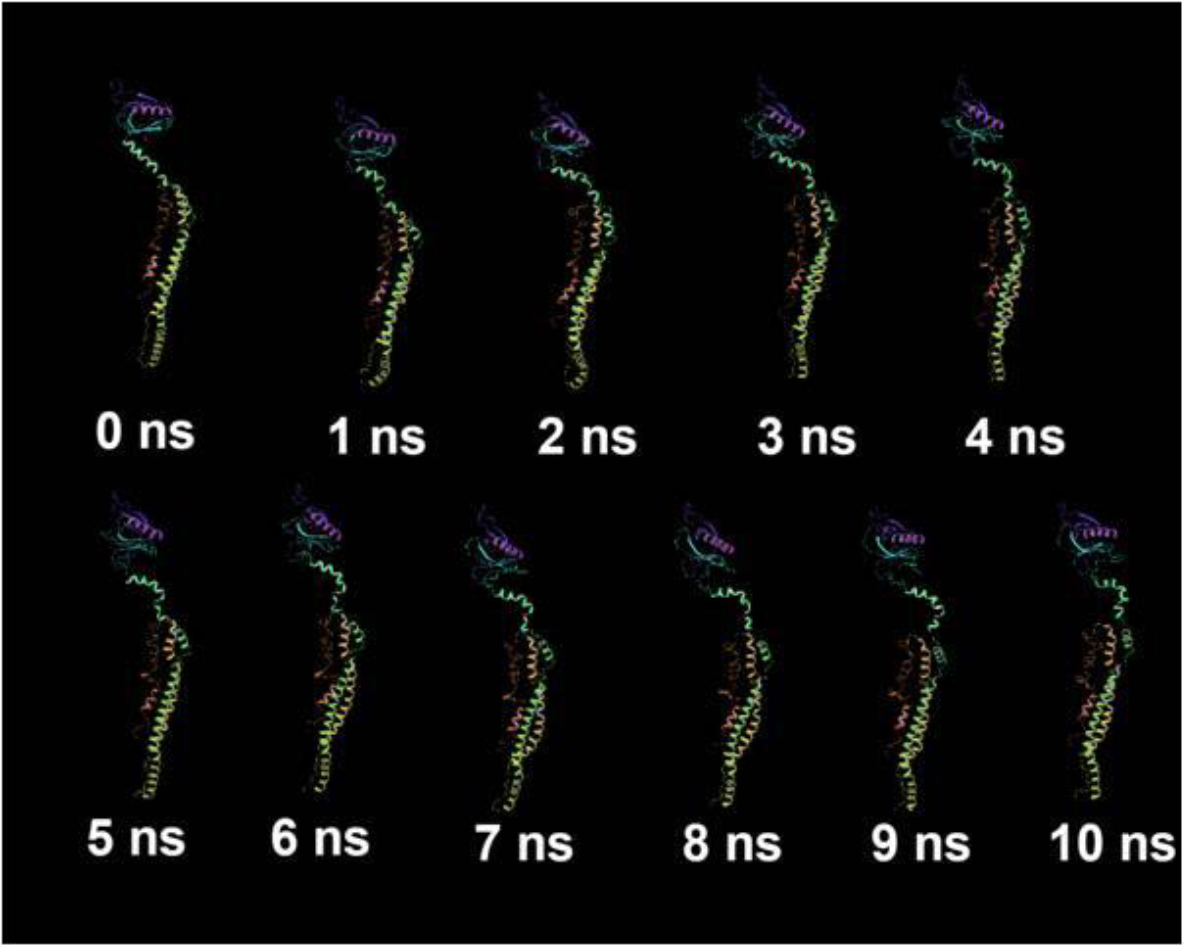
Snapshot of the rs3816281 PLEK mutant trajectory for every nanosecond of the MD simulation.

**Fig 10:**
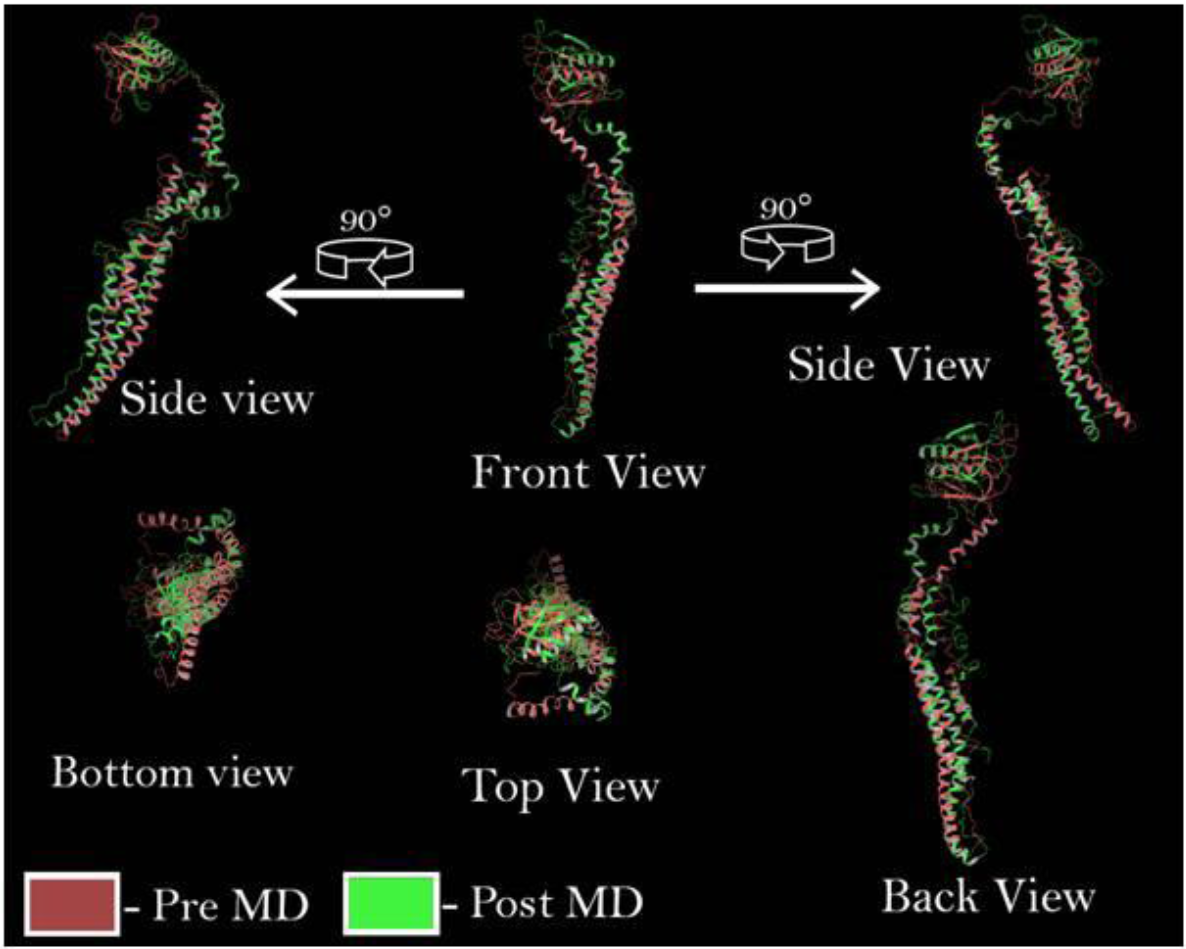
Visualization of the rs3816281 mutant PLEK protein superimposed of pre (Red) and post (Green) MD structures

## Discussion

Our study offers insight into the impact of SNP on the structure and function of PLEK gene and enables an insight on the molecular evolution of the PLEK gene.

Using insilico methods we have shown that the human PLEK gene has evolved in between Silurian to Devonian times and were subjected to selection pressures during this time (Fig. 1) while in the Jurassic and Cretaceous times, the PLEK gene has undergone changes indicating a process of stabilization and bearing correlation with the mammalian radiation (Fig. 1). This study finds importance in understanding and tracing the changes in the PLEK gene across evolutionary time spans.

From this study we report two damaging SNP’s: rs17035364 and rs3816281 in the PLEK gene which causes structure function alteration and leads to an impairment in the biological function.The RMSD, RMSF, total energy, radius of gyration and intramolecular hydrogen bonds analysis performed on the mutant and wild type/native PLEK proteins reveal the plausible malfunction of the protein due to structural destabilization. Two SNPs (rs3816281 and rs17035364) revealed of damaging consequences confirmed by MD simulation analysis results (Table1) as compared to the wild type or native proteins. This was also confirmed by RMSD (Fig. 2), RMSF (Fig. 3) values of native /wild types vs. mutant proteins revealing that the rs3816281 mutated protein has the highest degree of flexibility (Fig. 3).The total energy parameters obtained from the MDS trajectories of the native/wild type and mutant proteins revealed marked differences, being significantly higher for rs17035364 and significantly lower for rs3816281 as compared to the wild/native form (Fig. 4). The comparison on intramolecular hydrogen bonds revealed that rs3816281 has greater flexibility compared to the wild type/native or rs17035364 forms (Fig 5) which confirms the higher degree of flexibility from the RMSF data. The radius of gyration parameter allowed us to evaluate the native and mutant types based on their compactness.

rs17035364 mutation possessing protein revealed highest compactness with a mean radius of gyration of 46.05 A as compared to the wild type/native form while rs3816281 mutated protein revealed least compactness (Fig. 6)

We report observation of structural changes the SNPs at different time points. (Fig 7–10).The structural changes in the two SNPsrs17035364 and rs3816281 could be observed from the snapshot of the PLEK mutant trajectory for every nanosecond of the MD simulation (Fig. 7, 9) and superimposed images of pre (Red) and post (Green) MD structures (Fig. 8, 10).

Our study proves conclusively through insilico approaches that (i) changes in the PLEK gene has been through different evolutionary time points and that(ii) SNPs in the PLEK gene could cause structural and functional alterations in the PLEK proteins. We report rs17035364 and rs3816281 induced structure and function changes in the PLEK protein and these SNPs could be targeted for potential therapeutic purposes.

